# Cell type-dependent effects of ellagic acid on cellular metabolism

**DOI:** 10.1101/319533

**Authors:** Alexandra L. Boehning, Safia A. Essien, Erica L. Underwood, Pramod K. Dash, Darren Boehning

## Abstract

Ellagic acid is a botanical polyphenol which has been shown to have numerous effects on cellular function. Ellagic acid can induce apoptosis and inhibit the proliferation of various cancer cell types *in vitro* and *in vivo*. As such, ellagic acid has attracted significant interest as a potential chemotherapeutic compound. One mechanism by which ellagic acid has been proposed to affect cellular physiology is by regulating metabolic pathways. Here we show the dose-dependent effects of ellagic acid on cellular energy production and downstream induction of the apoptotic program in HEK293, HeLa, MCF7, and HepG2 cells. At physiologically relevant doses, ellagic acid has pleiotropic and cell-type specific effects on mitochondrial function. At high doses ellagic acid can also influence glycolytic pathways and induce cell death. Our results demonstrate that ellagic acid can influence mitochondrial function at therapeutically relevant concentrations. The observed effects of ellagic acid on cellular respiration are complex and cell type-specific, which may limit the chemotherapeutic utility of this compound.

## 1. INTRODUCTION

Ellagic acid (EA)^1^ is a polyphenolic compound enriched in nuts and berries. Like all polyphenols EA is an antioxidant [1, 2] and has many cellular effects including inhibiting cancer cell proliferation [3, 4]. Ellagic acid is bioavailable when taken orally and can briefly reach serum concentrations up to several hundred nanomolar in human subjects [5–7]. Metabolism of EA by microbial action in the gut leads to the production of urolithins which are also bioactive and achieve much higher serum concentrations with a longer half-life compared to EA [5]. Like EA, bioactive urolithin metabolites are antioxidants and have anti-cancer activity [5, 8] but may have unique cellular targets [9].

The number of cellular processes purported to be modulated by EA is impressive [4]. In non-transformed cells, EA is protective against oxidative insults [10–12]. In contrast, EA has antiproliferative and apoptotic effects in cancer cells. These seemingly contradictory activities of EA may be due to the unique energetic demands of the tumor microenvironment. Transformed cells require high levels of ATP, NADPH, and cellular building blocks such amino acids to support cell division and migration, often in the context of a hypoxic environment. In cancer cells EA appears to target these unique metabolic demands by regulating mitochondrial function. Ellagic acid causes mitochondrial depolarization in pancreatic, B cell, neuroblastoma, and bladder cancers [13–16], in most cases indirectly by activating apoptotic pathways. In B cell chronic lymphocytic leukemia EA has direct effects on mitochondria resulting in ROS production and release of pro-apoptotic factors from mitochondria [13]. There is also evidence that EA can affect cellular ATP production by targeting glycolysis. Ellagic acid downregulates the expression of the sodium/hydrogen exchanger 1 (NHE1) leading to cellular acidification and inhibition of glycolytic flux in endometrial cancer cells [17]. In T cell lymphoma EA can inhibit the activity of lactate dehydrogenase to decrease glycolysis [18]. One important caveat regarding many *in vitro* studies investigating EA is that the experimental effects on cellular physiology are not apparent at EA concentrations which are attainable in human serum (less than 500nM [3, 8, 19]).

In this study, we investigated the dose-dependent effects of EA on cellular ATP production in one non-transformed cell line (HEK293) and three commonly used cancer cell lines derived from different primary tumors (HeLa, MCF7, and HEPG2). Total ATP levels and *in situ* mitochondrial function in living cells were determined to evaluate the generality of the effects of EA on cellular metabolism at physiologically relevant concentrations. Finally the ability of EA to induce apoptotic cell death was investigated.

## 2. MATERIALS AND METHODS

### 2.1. Cell culture and materials

HepG2 human hepatocellular carcinoma, MCF7 human breast adenocarcinoma, HeLa human cervical adenocarcinoma, and HEK293 human embryonic kidney epithelial cells were purchased from ATCC. HepG2, HeLa, and HEK293 cells were cultured in Dulbecco’s Modified Eagle Medium (DMEM) supplemented with 10% fetal bovine serum. MCF7 cells were cultured in minimal essential medium (MEM) supplemented with 0.01mg/ml insulin and 10% fetal bovine serum. Cell lines were authenticated by ATCC by STR analysis and were certified mycoplasma free. Culture media and fetal bovine serum were purchased from Gibco/ThermoFischer Scientific. Extracellular flux reagents were purchased from Agilent Technologies. JC-1 was purchased from Molecular Probes/ThermoFischer Scientific. All other materials were purchased at the highest purity available from Sigma-Aldrich.

### 2.2. Cellular ATP levels by endpoint assay

Total ATP levels using the endpoint assay in Figure 1 were quantified using the CellTiter-Glo luminescent kit (Promega) in 96 well format. Three biological replicates averaged from eight technical replicates were used for each data point. Where noted, addition of 5 mM 2-deoxy-D-glucose (2DG) was concurrent with ellagic acid (or vehicle), and the cells were lysed for the endpoint assay 24 hours later. Luciferase activity was measured using a microplate reader essentially as described by the manufacturer. Glycolytic capacity was calculated as the difference between ATP in the presence and absence of 2DG normalized to vehicle. Respiratory capacity was simply ATP levels in the presence of 2DG normalized to vehicle. An unpaired t test was performed to compare vehicle to different concentrations of ellagic acid for each parameter quantified. For all experiments in this paper a statistical comparison was considered significant compared to vehicle control at p < 0.05 and is indicated on the figures with an asterisk. A two-tailed Student’s t test was used for all comparisons between two groups. Data in all figures are presented as the mean +/− standard error of the mean. Investigators were not blinded to the treatment conditions.

**Figure 1.**
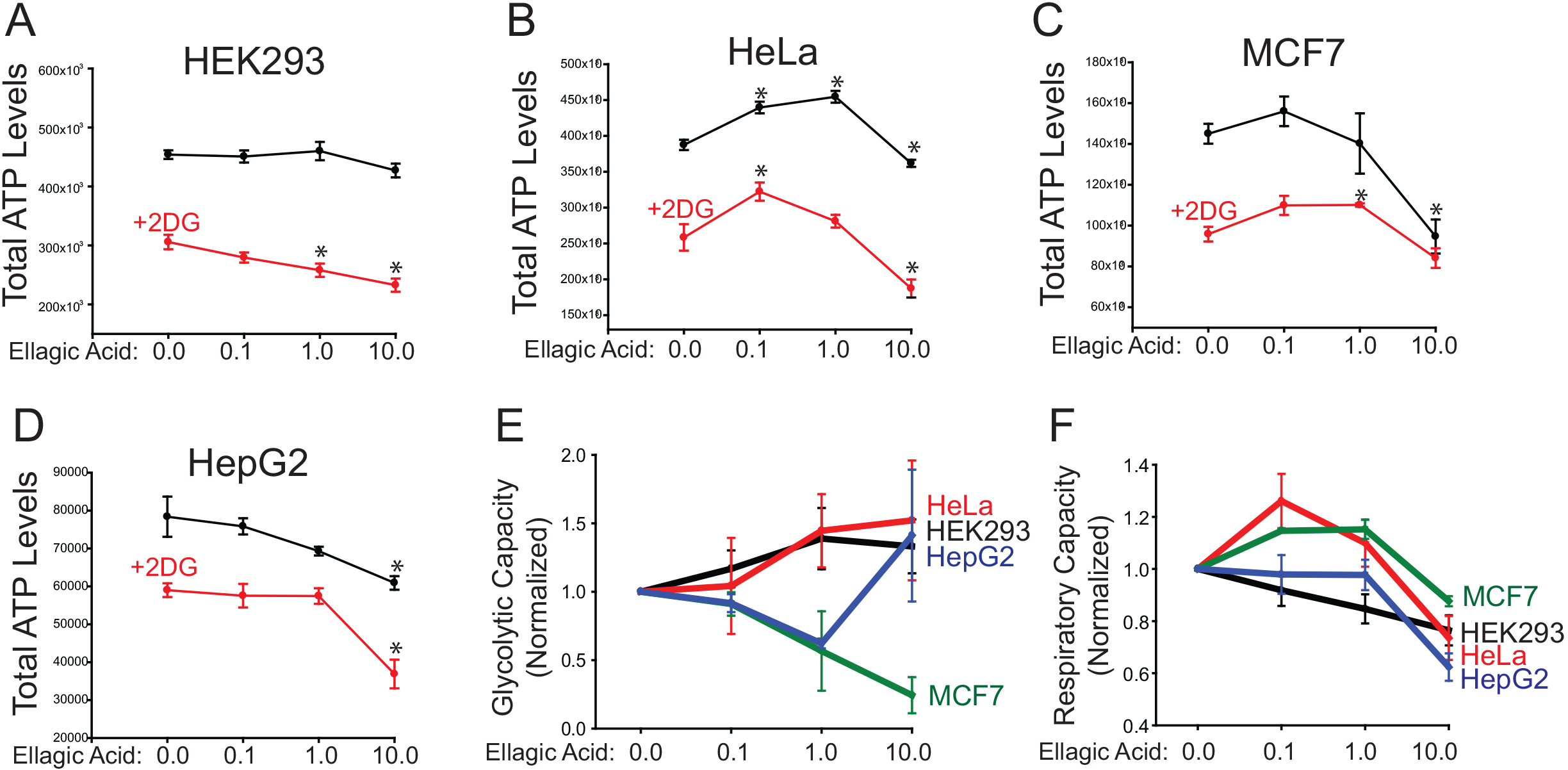
Ellagic acid impacts cellular ATP levels. A) Total HEK293 cellular ATP levels in the presence and absence of the glycolysis inhibitor 2-deoxy-D-glucose (2DG) after 24 hours treatment with the indicated concentrations of ellagic acid (in μM). The data are pooled from three separate experiments. The same experiment was repeated in HeLa (B), MCF7 (C), and HepG2 (D) cells. Data points represent the mean +/− s.e.m. of three separate determinations. *p < 0.05 versus control (vehicle) using an unpaired t test. E) Normalized glycolytic capacity of each cell type. F) Normalized respiratory capacity of each cell type.

### 2.3. Extracellular flux measurements

Cellular oxygen consumption rate was calculated in 96 well format using the Seahorse XF Cell Mito Stress Kit (Agilent Technologies). Assays were set up essentially as described by the manufacturer. The night before the assay, each cell type was plated at a density of 20,000 cells per well. All cells were treated the next day with ellagic acid in octuplet 1 hour prior to measurement of OCR rate. The concentration of FCCP uncoupler was varied based upon the cell type: 0.5μM final in HeLa, HepG2 and HEK293 assays; 0.25μM final in the MCF7 assay. Oxygen consumption rates were measured in a Seahorse XF Analyzer. The following parameters were quantified post-acquisition using formulas provided by the manufacturer: basal respiration, non-mitochondrial consumption, maximal respiration, proton leak, ATP production, and spare respiratory capacity from data pooled from three separate biological replicates. All values are reported as oxygen consumption rate (OCR), with units of pmol/min. Values on the outer edges of the plate were omitted. Also eliminated were any values reporting a negative OCR. An unpaired t test was performed to compare vehicle to different concentrations of ellagic acid for each parameter quantified.

### 2.4. JC-1 staining and analysis

Cells were passed onto coverslips 24 hours prior to imaging. Cell loading and imaging was performed in imaging buffer composed of 107 mM NaCl, 20 mM HEPES, 2.5 mM MgCl_2_, 7.25 mM KCl, 11.5 mM glucose, 1 mM CaCl_2_, and 1% bovine serum albumin. The JC-1 dye was purchased from Thermofisher. The JC-1 dye was reconstituted in DMSO to a final concentration of 20 mg/ml and stored at −80 until use. Loading solution was prepared by diluting this stock to 50 μg/ml in imaging buffer. This solution was then vortexed extensively to facilitate dispersion. Finally, the solution was briefly sonicated with a tip sonicator and centrifuged at 5000 xg for 15 minutes to pellet any insoluble material. The supernatant was used for staining. Cells were loaded with JC-1 for 10 minutes at 37°C and the staining solution was replaced with imaging buffer until mounting on an inverted live cell imaging microscope. Simultaneous imaging of both monomer and aggregate were accomplished by excitation at 480 nm and measuring dual emission at 525 nm (monomer) and 620 nm (aggregate). The multi-dichroic beamsplitter was Chroma Technology part number 86007bs. Fluorescent mages were acquired every three seconds on a Nikon TiS inverted microscope with a 40X oil immersion Nikon Super Flour objective with a numerical aperture of 1.30 and a working distance of 0.22mm. Images were acquired with a Photometrics Evolve deep cooled, back thinned EMCCD using Metafluor software. Each cell type was imaged as a single field in biological triplicates, and the total number of single cells analyzed is listed directly on the traces in Figure 4. Image analysis was done using ImageJ. Mitochondrial membrane potential was inferred from the standard deviation of the red channel fluorescence from a region of interest encompassing an entire single cell. The rationale behind this approach is that highly polarized mitochondria would have fluorescent red puncta with a high average standard deviation on average over the cell area, whereas depolarized mitochondria would cause monomerization and loss of red puncta and a subsequent decrease in standard deviation. A similar approach has been used extensively to monitor cytochrome C-GFP fusion protein release from mitochondria [20, 21]. This approach also facilitated comparisons between the cell lines as those cells which had green channel fluorescence barely above background (such as HeLa) had artificially inflated red/green ratios.

### 2.5. Caspase-3 activity

Caspase-3 and caspase-6 enzymatic activities were performed exactly as described previously 24 hours after treatment with ellagic acid [20]. Three biological replicates averaged from three technical replicates were used for each data point. A t test was performed to compare vehicle to different concentrations of ellagic acid for each parameter quantified.

## 3. RESULTS

### 3.1. Ellagic acid inhibits ATP production

We first investigated the effects of EA on total ATP levels in four commonly used cell lines: human embryonic kidney HEK293 cells, cervical carcinoma HeLa cells, metastatic breast cancer MCF7 cells, and hepatocellular carcinoma HepG2 cells. Each cell line was incubated with either vehicle (DMSO) or 0.1, 1.0, or 10 μM EA for 24 hours. The cells were then lysed and total ATP levels were quantified using a luciferase-based assay [22]. At all doses EA had no effect on ATP levels in HEK293 cells (Figure 1A). HeLa cells had a complex response to EA, with increases at 0.1 and 1.0 μM, and a slight reduction at 10 μM compared to control (Figure 1B). Both MCF7 and HepG2 cells had no differences in ATP concentration when treated with 0.1 μM and 1.0 μM EA, and a decrease at 10 μM EA. To investigate whether EA was affecting glycolysis or respiration, we measured ATP levels in the presence of the glycolysis inhibitor 2-deoxy-D-glucose (2DG; red plots in Figure 1A-D). As expected, 2DG reduced cellular ATP levels in all cell types. In HEK293, HeLa and HepG2 cells, EA further reduced ATP levels in the presence of 2DG (Figure 1A, C-D). In contrast to the other three cell types, in MCF7 cells there was no reduction in ATP levels in the presence of 2DG (Figure 1C). To determine the glycolytic capacity at each EA concentration, we subtracted total ATP levels from the levels measured in the presence of 2DG and normalized the data to control (vehicle). As shown in Figure 1E, EA only reduced glycolysis in MCF7 cells. Normalized respiratory capacity was reduced in all cells types except MCF7 cells using this assay consistent with the large body of literature indicating EA can affect mitochondrial function. One important caveat to these findings is that 2DG uptake and retention is dependent upon glucose transporter and hexokinase activity respectively. It is well known that cancer cells upregulate both of these enzymatic activities to different extents [23, 24], thus some of these cell type-specific differences may relate to alterations in glycolytic pathways. It is also not ideal to use an endpoint assay to measure ATP production. We therefore tested more directly the effects of EA on cellular respiration.

### 3.2. Ellagic acid has cell-type specific effects on respiration

To investigate the acute effects of EA on mitochondrial function *in situ* in living cells, we measured the oxygen consumption rate (OCR) in the presence of various inhibitors to evaluate six key parameters of mitochondrial function: basal respiration, maximal respiration, ATP production, spare respiratory capacity, proton leak, and non-mitochondrial oxygen consumption using the Seahorse extracellular flux assay [25, 26]. For these experiments each cell type was treated with various concentration of EA for 1 hour prior to measuring the oxygen consumption rate. Treatment of HEK293 cells with EA resulted in modest increases in basal respiration, maximal respiration, ATP production and spare respiratory capacity at a physiologically relevant dose (100 nM; Figure 2A). Higher doses did not have any effect with the exception of maximal respiratory rates. The complex dose-dependent effects on OCR is consistent with previous studies investigating mitochondrially-targeted antioxidants [27]. HeLa cells also had increased basal respiration and ATP production at low EA doses, however there were also negative effects on spare respiratory capacity and proton leak (Figure 2B). In contrast to HEK293 and HeLa cells, 100 nM EA decreased basal respiration and ATP production and increased the spare respiratory capacity in MCF7 cells (Figure 2C). In all three cell types, there was no clear dose dependent effects of EA. However, the observed effects on OCR occurred at physiologically relevant concentrations of EA (100 nM). HepG2 cells only had changes in OCR at the highest dose of EA (10 μM; Figure 2D), and resulted in decreased basal respiration, maximal respiration, spare respiratory capacity, proton leak, and non-mitochondrial oxygen consumption. To facilitate visualization of the broad and dose-dependent effects of EA on mitochondrial function, the oxygen consumption rate data were combined onto a single graph for each cell type in Figure 4. Replotting in this manner was also more useful to determine the effect sizes of EA on OCR. In HEK293 cells there was clear dose-dependent effects on maximal respiration with most other parameters remaining unchanged (Figure 3A). In both HeLa and MCF7 cells, EA had very modest effects on mitochondrial OCR with no obvious patterns to its action (Figure 3B-C). In contrast to the other three cells types, EA had robust and dose-dependent effects on OCR in HepG2 cells (Figure 3D), with nearly all parameters inhibited after treatment with 10 μM EA. This finding correlates well with our measurements of endpoint ATP levels in the presence of 2DG to inhibit glycolysis, where EA had the largest effect on cellular ATP levels in HepG2 cells (Figure 1D). It can be concluded that the effects of acute treatment with EA on mitochondrial respiration are complex and highly cell-type specific. This may be related to differences in mitochondrial content, morphology, and ultimately function in these four cell types. To evaluate these possibilities, we next performed single cell imaging of mitochondrial polarization after acute treatment with EA.

**Figure 2.**
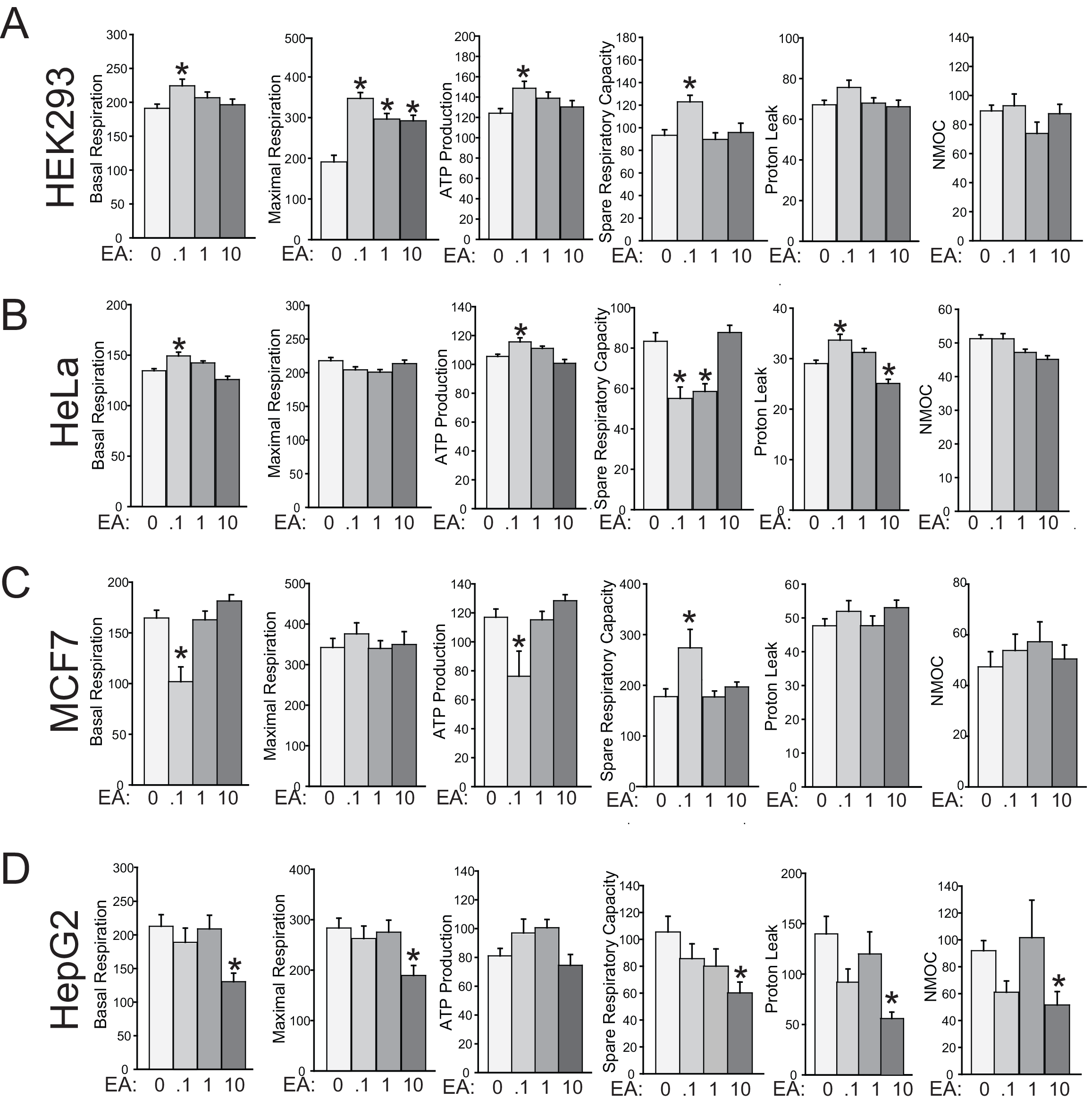
Dose-dependent effects of ellagic acid on oxygen consumption rate. A) Basal respiration, maximal respiration, ATP production, spare respiratory capacity, proton leak, and non-mitochondrial oxygen consumption (NMOC) in HEK293 cells determined by extracellular flux measurements at various ellagic acid (EA) concentrations (in μM). Ellagic acid was added 1 hour prior to measurement. The data are averages of the oxygen consumption rates (in pmol/min) and are pooled from three separate experiments. The same experiment was repeated in HeLa (B), MCF7 (C), and HepG2 (D) cells. Data points represent the mean +/− s.e.m. of three separate determinations. *p < 0.05 versus control (vehicle) using an unpaired t test.

**Figure 3.**
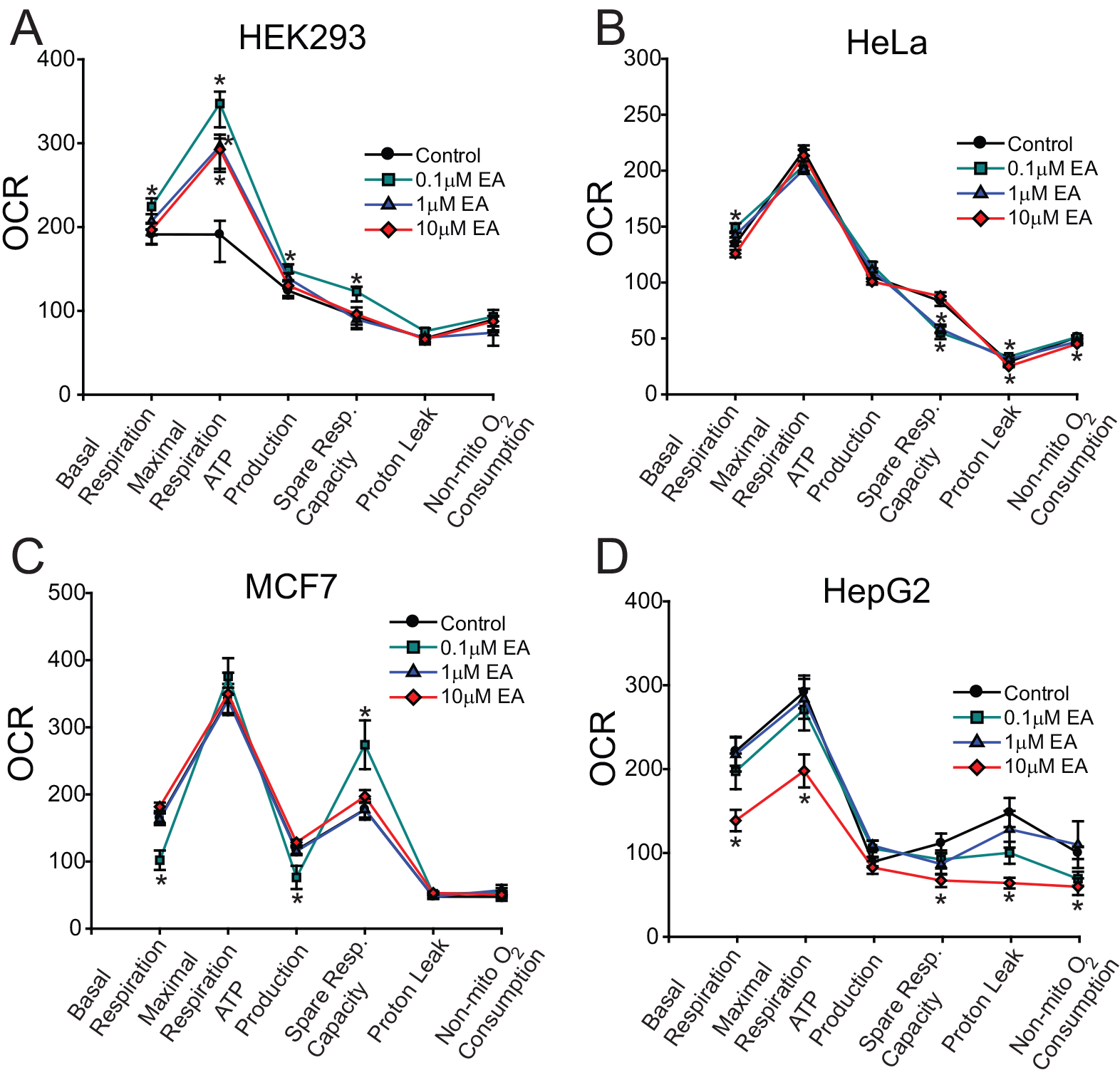
Ellagic acid has cell-type specific effects on mitochondrial function. A) Pooled respiratory function data for HEK293 cells at various ellagic acid (EA) concentrations. Data points represent the mean +/− s.e.m. of the oxygen consumption rate (OCR) of three separate determinations. *p < 0.05 versus control (vehicle) using an unpaired t test. The same experiment was repeated in HeLa (B), MCF7 (C), and HepG2 (D) cells.

**Figure 4.**
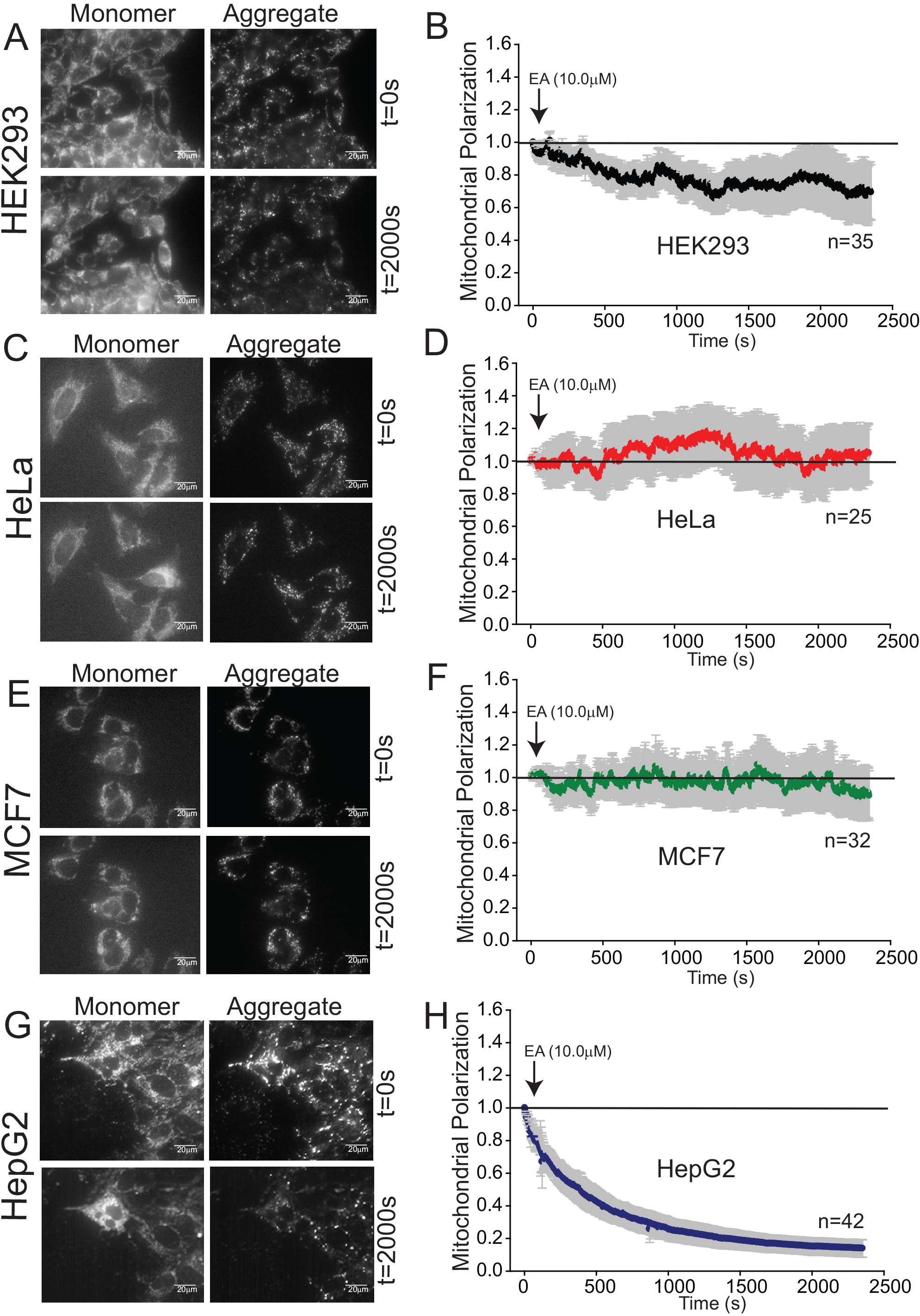
Effects of acute administration of ellagic acid on mitochondrial membrane potential. A) Representative JC-1 images of the green fluorescent monomer and red fluorescent aggregate at time zero and 2000 seconds after 10 μM ellagic acid (EA) addition in HEK293 cells. B) Normalized mitochondrial polarization (as determined by red aggregate fluorescence) before and after EA addition in HEK293 cells. Total number of single cells analyzed is indicated. Data points represent the mean +/− s.e.m. of three separate determinations. 0,D) Same experiment in HeLa cells. Resting green fluorescence (monomer) is very low in this cell line. E,F) Same experiment in M0F7 cells. G,H) Same experiment in HepG2 cells. In all graphs a solid line represents no change from time zero. Error bars are colored grey.

### 3.3. Ellagic acid has cell-type specific effects on mitochondrial polarization

Several studies have shown that EA induces mitochondrial depolarization [13–16]. Most studies use chronic EA treatment models which also induce cell death signaling pathways which are known to depolarize mitochondria. Thus, it is difficult to ascertain from these studies whether the effects of EA in mitochondrial function were direct or indirect. To investigate the acute effects of EA on mitochondrial polarization, we employed real time *in situ* imaging of live cells using single-cell JC-1 imaging. The dye JC-1 aggregates and fluoresces red in polarized mitochondria, while the green monomeric form predominates in depolarized mitochondria [28, 29]. Often a single mitochondria will have significant heterogeneity in membrane potential and will have regions where red and green fluorescence predominates [28]. Staining of HEK293 cells with JC-1 reveals clear mitochondrial localization with red/green heterogeneity (Figure 4A). Treatment with 10 μM EA causes a slight reduction in mitochondrial polarization over a 40 minute time course (Figure 4B). Ellagic acid had no effects on mitochondrial polarization in HeLa or MCF7 after 40 minutes of treatment (Figure 4C-F). In contrast, EA treatment of HepG2 cells caused a dramatic and almost immediate reduction in mitochondrial polarization (Figure 4G-H). This finding likely explains the broad effects of EA on mitochondrial respiration observed in HepG2 cells determined by extracellular flux analysis (Figures 2–3). In all cell types there was no obvious changes in mitochondrial morphology after EA treatment as determined by the green JC-1 fluorescence (Figure 4 A,C,E,G). We conclude that the effects on EA on ATP production and mitochondrial respiration are remarkably cell type specific. Importantly, these findings also highlight that potential therapeutics which have an effect on mitochondrial function in one of these cell types may not be generalizable to other cell types.

### 3.4. Ellagic acid is not a robust inducer of caspase activation

Ellagic acid has been shown to specifically induce apoptosis in cancer cells [13, 14, 18, 30–34] while having cytoprotective activity in non-transformed cells [2, 4, 10, 12, 19, 35]. To determine whether EA activated apoptotic pathways, we measured caspase-3 catalytic activities in cells treated with EA for 24 hours. Treatment of non-transformed HEK293 cells with EA caused a slight reduction in basal caspase-3 activity (Figure 5A). In HeLa cells we found that a supraphysiologic dose of EA (10 μM) could increase caspase-3 catalytic activity about 2 fold (Figure 5B). For comparison, a robust apoptotic inducer of apoptosis in this cell line such as staurosporine will cause a 10-50 fold increase in caspase-3 activity [20, 36]. Ellagic acid had no statistically significant effect on caspase 3 activity in HepG2 cells (Figure 5C) despite strong effects on mitochondrial function in this cell type (Figures 2–4). MCF7 cells have a mutation in the caspase-3 gene which eliminates the expression of the pro-enzyme [37]. We therefore measured caspase-6 activity in this cell line. We could not detect any caspase-6 activity in MCF7 cells, even under basal conditions. Thus, at least in these four cell lines, EA is not a robust inducer of the apoptotic program. These findings also suggest that the changes in glycolytic flux or OCR observed in Figures 1–4 cannot be explained by activation of cell death pathways.

**Figure 5.**
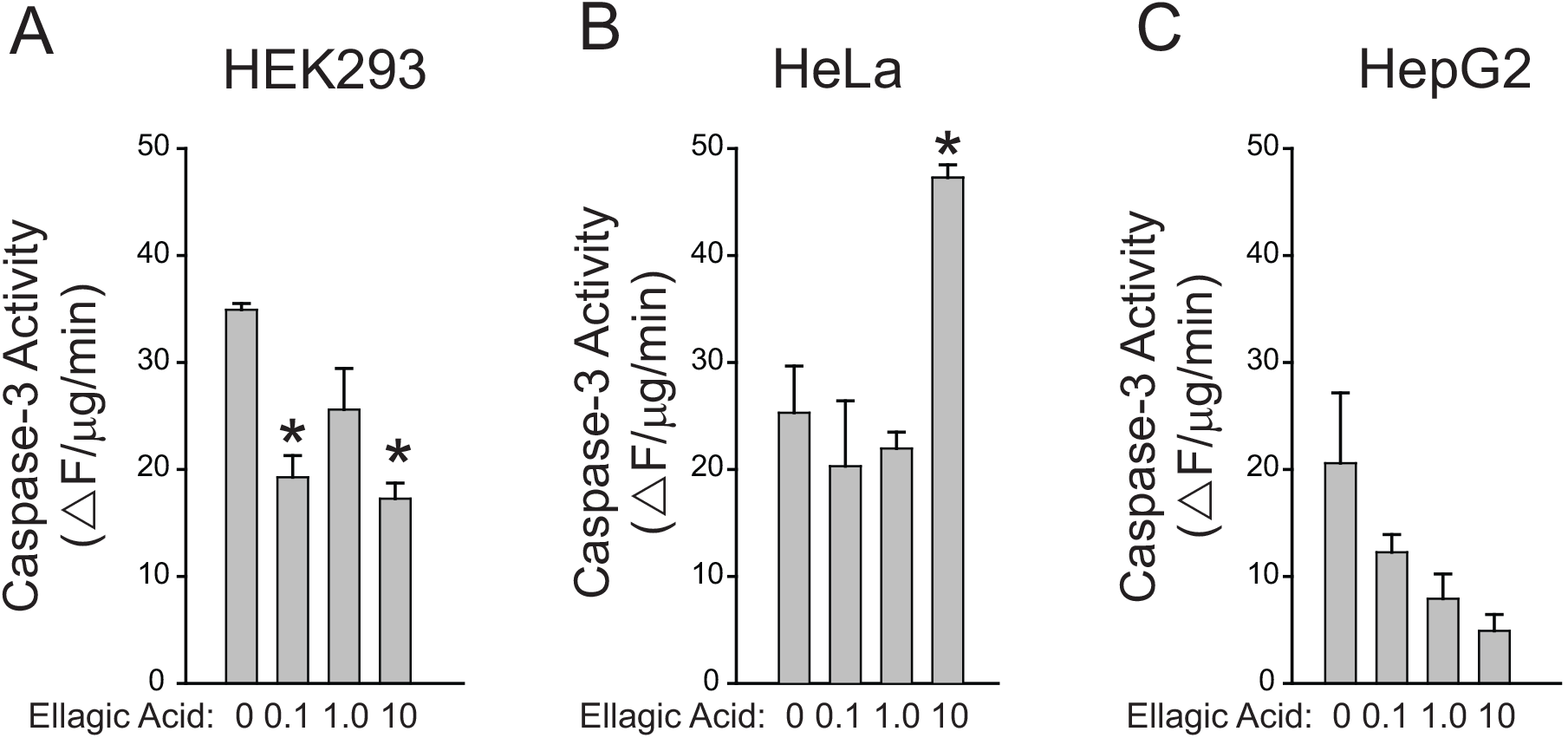
Ellagic acid is not a potent regulator of caspase activity. A) Caspase-3 enzymatic activity in HEK293 cells after treatment with the indicated concentrations of ellagic acid for 24 hours. Caspase-3 enzymatic activity was measured using a fluorogenic caspase-3 substrate on cell lysates (Boehning et al., 2003). Data points represent the mean +/− s.e.m. of three separate determinations. The units are change in fluorescent intensity per μg per minute. *p < 0.05 versus control (vehicle) using an unpaired t test. The same experiment was repeated in HeLa (B), and HepG2 (C) cells. As noted in the text MCF-7 cells have a mutation at the CASP3 locus eliminating expression of this enzyme. Using an identical enzyme assay with a fluorogenic caspase-6 substrate resulted in no detectable caspase-6 activity in control or EA treated cells and as such is not graphed.

## 4. DISCUSSION

This study demonstrates that EA has highly cell type-dependent effects on cellular metabolism. Within individual cell types, the multitude of effects on mitochondrial function suggest that there is not a single target of action for this compound. Not surprisingly, some of the effects of EA mimic the action of antioxidants [4, 27]. Almost all previous studies using EA have utilized doses which are not achievable in human serum. In this study we see modest effects on mitochondrial function at physiologically relevant doses (100 nM). However, our findings also highlight that the effects of EA are cell type specific. This may limit the usefulness of this compound as a therapeutic for cancer. Urolithin metabolites of EA achieve much higher serum levels and have a longer half-life [8, 38]. It will be useful in future studies to examine the role of these compounds on mitochondrial function.

A major question remains as to why there are differential responses in these cell lines. Ellagic acid can be transported into the cell *via* several mechanisms. Organic anion transporters of the *SLCO* (OATP) and *SLC22A* (OAT1 and OAT4) families can transport EA and facilitate intracellular accumulation [39–41]. In addition, there is evidence that EA can be transported by the sodium-dependent glucose transporter SGLT1 [41, 42]. It is well known that organic anion transporters and sodium-dependent glucose transporters are differentially expressed in cancer cells and contribute to cancer development [43–46]. These transport mechanisms are dependent upon ATP and sodium levels, both of which can be altered in cancer [47, 48]. Thus, our results might ultimately be explained by differential EA uptake rates and accumulation by each cell type. It is also clear from our data that there are differences in baseline mitochondrial function (Figures 2–4) and morphology (Figure 4) between the four cell types which may explain the differential responses to EA treatment. Future studies will be focused on determining the precise molecular target(s) of action.

## DISCLOSURE OF INTEREST

The authors declare that they have no conflicts of interest concerning this article.

## ACKNOWLEDGEMENTS

We would like to thank Tara Fischer for very useful discussions and guidance regarding the Seahorse XF assay.

## FUNDING SOURCES

This work was support by National Institute of Health National Institute of General Medical Sciences [Grant GM081685] and the National Institute of Neurological Disorders and Stroke [Grants NS090935, NS087149]

## Footnotes

1 **Non-standard abbreviations:** 2DG: 2-Deoxy-D-glucose; EA: Ellagic Acid; OCR: Oxygen consumption rate

## REFERENCES

[1] F. Di Meo, V. Lemaur, J. Cornil, R. Lazzaroni, J.L. Duroux, Y. Olivier, P. Trouillas, Free radical scavenging by natural polyphenols: atom versus electron transfer, J Phys Chem A 117(10) (2013) 2082–92.

[2] K.I. Priyadarsini, S.M. Khopde, S.S. Kumar, H. Mohan, Free radical studies of ellagic acid, a natural phenolic antioxidant, J Agric Food Chem 50(7) (2002) 2200–6.

[3] H.M. Zhang, L. Zhao, H. Li, H. Xu, W.W. Chen, L. Tao, Research progress on the anticarcinogenic actions and mechanisms of ellagic acid, Cancer Biol Med 11(2) (2014) 92–100.

[4] W.R. Garcia-Nino, C. Zazueta, Ellagic acid: Pharmacological activities and molecular mechanisms involved in liver protection, Pharmacol Res 97 (2015) 84–103.

[5] I. Kang, T. Buckner, N.F. Shay, L. Gu, S. Chung, Improvements in Metabolic Health with Consumption of Ellagic Acid and Subsequent Conversion into Urolithins: Evidence and Mechanisms, Adv Nutr 7(5) (2016) 961–72.

[6] A. González-Sarrías, R. García-Villalba,M.A. Núñez-Sánchez, J. Tomé-Carneiro, P. Zafrilla, J. Mulero, F.A. Tomás-Barberán, J.C. Espín, Identifying the limits for ellagic acid bioavailability: A crossover pharmacokinetic study in healthy volunteers after consumption of pomegranate extracts, J Funct Foods 19(A) (2015) 225–235.

[7] N.P. Seeram, R. Lee, D. Heber, Bioavailability of ellagic acid in human plasma after consumption of ellagitannins from pomegranate (Punica granatum L.) juice, Clin Chim Acta 348(1-2) (2004) 63–8.

[8] J.C. Espin, M. Larrosa, M.T. Garcia-Conesa, F. Tomas-Barberan, Biological significance of urolithins, the gut microbial ellagic Acid-derived metabolites: the evidence so far, Evid Based Complement Alternat Med 2013 (2013) 270418.

[9] R. Vicinanza, Y. Zhang, S.M. Henning, D. Heber, Pomegranate Juice Metabolites, Ellagic Acid and Urolithin A, Synergistically Inhibit Androgen-Independent Prostate Cancer Cell Growth via Distinct Effects on Cell Cycle Control and Apoptosis, Evid Based Complement Alternat Med 2013 (2013) 247504.

[10] F. Firdaus, M.F. Zafeer, M. Waseem, E. Anis, M.M. Hossain, M. Afzal, Ellagic acid mitigates arsenic-trioxide-induced mitochondrial dysfunction and cytotoxicity in SH-SY5Y cells, J Biochem Mol Toxicol 32(2) (2018).

[11] A. Dhingra, R. Jayas, P. Afshar, M. Guberman, G. Maddaford, J. Gerstein, B. Lieberman, H. Nepon, V. Margulets, R. Dhingra, L.A. Kirshenbaum, Ellagic acid antagonizes Bnip3-mediated mitochondrial injury and necrotic cell death of cardiac myocytes, Free Radic Biol Med 112 (2017) 411–422.

[12] M.R. Sepand, M.H. Ghahremani, K. Razavi-Azarkhiavi, M. Aghsami, J. Rajabi, H. Keshavarz-Bahaghighat, M. Soodi, Ellagic acid confers protection against gentamicin-induced oxidative damage, mitochondrial dysfunction and apoptosis-related nephrotoxicity, J Pharm Pharmacol 68(9) (2016) 1222–32.

[13] A. Salimi, M.H. Roudkenar, L. Sadeghi, A. Mohseni, E. Seydi, N. Pirahmadi, J. Pourahmad, Ellagic acid, a polyphenolic compound, selectively induces ROS-mediated apoptosis in cancerous B-lymphocytes of CLL patients by directly targeting mitochondria, Redox Biol 6 (2015) 461–71.

[14] C.C. Ho, A.C. Huang, C.S. Yu, J.C. Lien, S.H. Wu, Y.P. Huang, H.Y. Huang, J.H. Kuo, W.Y. Liao, J.S. Yang, P.Y. Chen, J.G. Chung, Ellagic acid induces apoptosis in TSGH8301 human bladder cancer cells through the endoplasmic reticulum stress- and mitochondria-dependent signaling pathways, Environ Toxicol 29(11) (2014) 1262–74.

[15] M. Edderkaoui, I. Odinokova, I. Ohno, I. Gukovsky, V.L. Go, S.J. Pandol, A.S. Gukovskaya, Ellagic acid induces apoptosis through inhibition of nuclear factor kappa B in pancreatic cancer cells, World J Gastroenterol 14(23) (2008) 3672–80.

[16] C.F. Alfredsson, M. Ding, Q.L. Liang, B.E. Sundstrom, E. Nanberg, Ellagic acid induces a dose-and time-dependent depolarization of mitochondria and activation of caspase-9 and -3 in human neuroblastoma cells, Biomed Pharmacother 68(1) (2014) 129–35.

[17] K.N.M. Abdelazeem, Y. Singh, F. Lang, M.S. Salker, Negative Effect of Ellagic Acid on Cytosolic pH Regulation and Glycolytic Flux in Human Endometrial Cancer Cells, Cell Physiol Biochem 41(6) (2017) 2374–2382.

[18] S. Mishra, M. Vinayak, Role of ellagic acid in regulation of apoptosis by modulating novel and atypical PKC in lymphoma bearing mice, BMC Complement Altern Med 15 (2015) 281.

[19] A. Zeb, Ellagic acid in suppressing in vivo and in vitro oxidative stresses, Mol Cell Biochem (2018).

[20] D. Boehning, R.L. Patterson, L. Sedaghat, N.O. Glebova, T. Kurosaki, S.H. Snyder, Cytochrome c binds to inositol (1,4,5) trisphosphate receptors, amplifying calcium-dependent apoptosis, Nat Cell Biol 5(12) (2003) 1051–61.

[21] J.C. Goldstein, N.J. Waterhouse, P. Juin, G.I. Evan, D.R. Green, The coordinate release of cytochrome c during apoptosis is rapid, complete and kinetically invariant, Nat Cell Biol 2(3) (2000) 156–62.

[22] M.A. Turman, A. Mathews, A simple luciferase assay to measure atp levels in small numbers of cells using a fluorescent plate reader, In Vitro Cell Dev Biol Anim 32(1) (1996) 1–4.

[23] M.L. Macheda, S. Rogers, J.D. Best, Molecular and cellular regulation of glucose transporter (GLUT) proteins in cancer, J Cell Physiol 202(3) (2005) 654–62.

[24] S.P. Mathupala, Y.H. Ko, P.L. Pedersen, Hexokinase II: cancer’s double-edged sword acting as both facilitator and gatekeeper of malignancy when bound to mitochondria, Oncogene 25(34) (2006) 4777–86.

[25] M. Pelletier, L.K. Billingham, M. Ramaswamy, R.M. Siegel, Extracellular flux analysis to monitor glycolytic rates and mitochondrial oxygen consumption, Methods Enzymol 542 (2014) 125–49.

[26] R. Wang, S.J. Novick, J.B. Mangum, K. Queen, D.A. Ferrick, G.W. Rogers, J.B. Stimmel, The acute extracellular flux (XF) assay to assess compound effects on mitochondrial function, J Biomol Screen 20(3) (2015) 422–9.

[27] C. Reily, T. Mitchell, B.K. Chacko, G. Benavides, M.P. Murphy, V. Darley-Usmar, Mitochondrially targeted compounds and their impact on cellular bioenergetics, Redox Biol 1(1) (2013) 86–93.

[28] S.T. Smiley, M. Reers, C. Mottola-Hartshorn, M. Lin, A. Chen, T.W. Smith, G.D. Steele, Jr., L.B. Chen, Intracellular heterogeneity in mitochondrial membrane potentials revealed by a J-aggregate-forming lipophilic cation JC-1, Proc Natl Acad Sci U S A 88(9) (1991) 3671–5.

[29] M. Reers, T.W. Smith, L.B. Chen, J-aggregate formation of a carbocyanine as a quantitative fluorescent indicator of membrane potential, Biochemistry 30(18) (1991) 4480–6.

[30] H.S. Chen, M.H. Bai, T. Zhang, G.D. Li, M. Liu, Ellagic acid induces cell cycle arrest and apoptosis through TGF-beta/Smad3 signaling pathway in human breast cancer MCF-7 cells, Int J Oncol 46(4) (2015) 1730–8.

[31] Y. Hagiwara, T. Kasukabe, Y. Kaneko, N. Niitsu, J. Okabe-Kado, Ellagic acid, a natural polyphenolic compound, induces apoptosis and potentiates retinoic acid-induced differentiation of human leukemia HL-60 cells, Int J Hematol 92(1) (2010) 136–43.

[32] M. Larrosa, F.A. Tomas-Barberan, J.C. Espin, The dietary hydrolysable tannin punicalagin releases ellagic acid that induces apoptosis in human colon adenocarcinoma Caco-2 cells by using the mitochondrial pathway, J Nutr Biochem 17(9) (2006) 611–25.

[33] S. Mishra, M. Vinayak, Ellagic acid induces novel and atypical PKC isoforms and promotes caspase-3 dependent apoptosis by blocking energy metabolism, Nutr Cancer 66(4) (2014) 675–81.

[34] D. Wang, Q. Chen, B. Liu, Y. Li, Y. Tan, B. Yang, Ellagic acid inhibits proliferation and induces apoptosis in human glioblastoma cells, Acta Cir Bras 31(2) (2016) 143–9.

[35] R. Atta Ur, F.N. Ngounou, M.I. Choudhary, S. Malik, T. Makhmoor, E.A.M. Nur, S. Zareen, D. Lontsi, J.F. Ayafor, B.L. Sondengam, New antioxidant and antimicrobial ellagic acid derivatives from Pteleopsis hylodendron, Planta Med 67(4) (2001) 335–9.

[36] A.M. Akimzhanov, J.M. Barral, D. Boehning, Caspase 3 cleavage of the inositol 1,4,5-trisphosphate receptor does not contribute to apoptotic calcium release, Cell Calcium 53(2) (2013) 152–8.

[37] R.U. Janicke, M.L. Sprengart, M.R. Wati, A.G. Porter, Caspase-3 is required for DNA fragmentation and morphological changes associated with apoptosis, J Biol Chem 273(16) (1998) 9357–60.

[38] A. Gonzalez-Sarrias, J.A. Gimenez-Bastida, M.T. Garcia-Conesa, M.B. Gomez-Sanchez, N.V. Garcia-Talavera, A. Gil-Izquierdo, C. Sanchez-Alvarez, L.O. Fontana-Compiano, J.P. Morga-Egea, F.A. Pastor-Quirante, F. Martinez-Diaz, F.A. Tomas-Barberan, J.C. Espin, Occurrence of urolithins, gut microbiota ellagic acid metabolites and proliferation markers expression response in the human prostate gland upon consumption of walnuts and pomegranate juice, Mol Nutr Food Res 54(3) (2010) 311–22.

[39] A.C. Whitley, D.H. Sweet, T. Walle, The dietary polyphenol ellagic acid is a potent inhibitor of hOAT1, Drug Metab Dispos 33(8) (2005) 1097–100.

[40] A.C. Whitley, D.H. Sweet, T. Walle, Site-specific accumulation of the cancer preventive dietary polyphenol ellagic acid in epithelial cells of the aerodigestive tract, J Pharm Pharmacol 58(9) (2006) 1201–9.

[41] X. Mao, L.F. Wu, H.J. Zhao, W.Y. Liang, W.J. Chen, S.X. Han, Q. Qi, Y.P. Cui, S. Li, G.H. Yang, Y.Y. Shao, D. Zhu, R.F. Wang, Y. You, L.Z. Zhang, Transport of Corilagin, Gallic Acid, and Ellagic Acid from Fructus Phyllanthi Tannin Fraction in Caco-2 Cell Monolayers, Evid Based Complement Alternat Med 2016 (2016) 9205379.

[42] K. Johnston, P. Sharp, M. Clifford, L. Morgan, Dietary polyphenols decrease glucose uptake by human intestinal Caco-2 cells, FEBS Lett 579(7) (2005) 1653–7.

[43] A. Obaidat, M. Roth, B. Hagenbuch, The expression and function of organic anion transporting polypeptides in normal tissues and in cancer, Annu Rev Pharmacol Toxicol 52 (2012) 135–51.

[44] S.K. Nigam, K.T. Bush, G. Martovetsky, S.Y. Ahn, H.C. Liu, E. Richard, V. Bhatnagar, W. Wu, The organic anion transporter (OAT) family: a systems biology perspective, Physiol Rev 95(1) (2015) 83–123.

[45] M. Roth, A. Obaidat, B. Hagenbuch, OATPs, OATs and OCTs: the organic anion and cation transporters of the SLCO and SLC22A gene superfamilies, British journal of pharmacology 165(5) (2012) 1260–87.

[46] H. Koepsell, The Na(+)-D-glucose cotransporters SGLT1 and SGLT2 are targets for the treatment of diabetes and cancer, Pharmacol Ther 170 (2017) 148–165.

[47] J.L. Fiske, V.P. Fomin, M.L. Brown, R.L. Duncan, R.A. Sikes, Voltage-sensitive ion channels and cancer, Cancer Metastasis Rev 25(3) (2006) 493–500.

[48] R.J. DeBerardinis, N.S. Chandel, Fundamentals of cancer metabolism, Sci Adv 2(5) (2016) e1600200.

